# TMEM63A, associated with hypomyelinating leukodystrophies, is an evolutionarily conserved regulator of myelination

**DOI:** 10.1101/2024.12.27.630433

**Authors:** Julia Halford, Amanda J. Senatore, Sage Berryman, Antonio Muñoz, Destinee Semidey, Ryan A. Doan, Adam M. Coombs, Brandon Noimany, Katie Emberley, Ben Emery, Kelly R. Monk, Swetha E. Murthy

**Affiliations:** Vollum Institute, Oregon Health & Science University; Portland, Oregon; Jungers Center for Neurosciences Research, Oregon Health & Science University; Portland, Oregon

## Abstract

Infantile hypomyelinating leukodystrophy 19 (HLD19) is a rare genetic disorder where patients exhibit reduced myelin in central nervous system (CNS) white matter tracts and present with varied neurological symptoms. The causative gene *TMEM63A* encodes a mechanosensitive ion channel whose role in myelination has not been explored. Our study shows that TMEM63A is a major regulator of OL-driven myelination in the CNS. In mouse and zebrafish, *Tmem63a* inactivation led to early deficits in myelination, recapitulating the HLD19 phenotype. OL-specific conditional mouse knockouts of *Tmem63a* exhibited transient reductions in myelin, indicating that TMEM63A regulates myelination cell-autonomously. We show that TMEM63A is present at plasma membrane and on lysosomes and modulates myelin/myelin-associated protein production. Intriguingly, HLD19-associated *TMEM63A* variants from patients blocked trafficking to cell membrane. Together, our results reveal an ancient role for TMEM63A in fundamental aspects of myelination *in vivo* and highlight two exciting new models for the development of treatments for devastating hypomyelinating leukodystrophies.

## INTRODUCTION

Monoallelic mutations in *TMEM63A* cause transient infantile hypomyelinating leukodystrophy 19 (HLD19, OMIM #618688) ^1–6^. Like other genetic leukodystrophies, HLD19 is characterized by a severe deficit in central nervous system (CNS) myelin identified by serial MRIs early in life. HLD19 patients present clinically as young as nine months of age with nystagmus, ataxia, and delayed motor development, displaying hypomyelination in the corpus collosum and deep and subcortical white matter. Interestingly, myelin deficits in *TMEM63A*- associated leukodystrophy can resolve with increasing age. Near-normal myelin was found in follow-up scans acquired around 2-3 years after the initial assessment for all patients described in Yan 2019 ^1^. However, there is a range in disease severity and possible outcomes for patients carrying *TMEM63A* variants. Patients identified in subsequent case studies suffer from a more severe manifestation of the disorder ^2,4^. Two subjects from the more recent studies did not experience resolution of their hypomyelination, including additional deficits in spinal cord white matter, which were not reported for other patients. The most recently recognized *TMEM63A* variant causes adult-onset neurological impairment, further broadening the range of *TMEM63A*- associated phenotypes ^7^. Collectively, the phenotypes across the cohort of HLD19 patients imply that TMEM63A, a mechanosensitive channel that converts mechanical stimuli into biological signals ^8,9^, is an indispensable molecule in the regulation of CNS myelination. However, such a role has not yet been explicitly demonstrated, and the underlying mechanism remains unknown.

In the CNS, myelin is formed by oligodendrocytes (OLs), which extend multiple processes that contact and iteratively wrap an axon segment before compacting to form the myelin sheath. By coating axons with lipid-rich, high-resistance membrane, myelin allows axons to conduct nerve impulses with high speed and acuity, making it so critical to nervous system function that any defects in myelination can cause debilitating symptoms in diseases such as the discussed inherited leukodystrophies and multiple sclerosis (MS). Even so, our current understanding of how CNS myelination is regulated remains incomplete.

Many extrinsic and intrinsic molecular factors facilitate interactions between neurons and OL lineage cells (OLCs) to drive myelination, such as transcriptional and posttranscriptional mechanisms or axonal surface ligands, secreted molecules, and axonal activity. More recently, it is becoming apparent that the mechanical properties of the cellular environment also play an essential role in OL development and myelination ^10^. The most notable mechanistic contribution of mechanosensation to CNS myelination is the ability of OLs to select axons based on their diameter ^11^. During development, large diameter axons are preferentially myelinated, with smaller diameter axons myelinated later or not at all ^12–14^. Compellingly, OLs have even been shown to myelinate biologically inert engineered nanofibers with the same preference for larger caliber fibers seen *in vivo* ^15^. These observations indicate that mechanical properties of axons are sufficient to initiate and regulate myelin formation in the absence of additional cues such as secreted or cell surface ligands, and that OLs have an intrinsic ability to sense axonal diameter through the membrane stretch exerted at the end of OL processes when exploring and contacting the axon shaft, although the identity of the molecule(s) that transduce these mechanical cues are currently unknown.

While it is well-established that the senses of touch and hearing depend on the transduction of physical force by mechanosensitive ion channels, it is emerging that mechanosensors participate in a range of previously unanticipated biological processes ^16,17^. By acting as precise membrane tension sensors, mechanosensitive channels are crucial for fundamental cellular processes such as migration, proliferation, and differentiation ^18^.

TMEM63A belongs to a family of bona fide mechanosensitive cation channels called OSCA/TMEM63, with conserved function in plants (OSCAs) and animals (TMEM63) ^8,9,19^. In recent years the *in vivo* roles of individual TMEM63 family members have been an emerging topic. Fly *Tmem63* is involved in food texture- and hygro- sensation and fat body regulation ^20–22^. In mice, *Tmem63a* has been implicated in inflammation-induced pain and *Tmem63b* in outer hair cell function ^23,24^, and both channels together are linked to lung inflation-induced surfactant release ^25^. The hypomyelination observed in the HLD19 patient cohort strongly suggests that TMEM63A is likely a mechanosensor in the regulation of myelination. Here, we use a range of model systems to investigate whether the myelin deficits in HLD19 may result from cell-autonomous loss of TMEM63A function in oligodendrocytes.

## RESULTS

### HLD19-associated variants of *TMEM63A* fail to localize to the plasma membrane

Since the initial discovery of heterozygous *TMEM63A* variants in patients with HLD19 ^1^, new variants in patients with similar presentation have been identified ^2,4,7^ (**Fig. 1A**). To determine whether the novel HLD19-associated variants also led to loss-of-function (LOF), we overexpressed *TMEM63A* in HEK cells and investigated channel function by stretching the membrane to apply a mechanical stimulus. We used PIEZO1-knockout HEK 293T (HEK-P1KO) cells, which otherwise lack mechanically induced currents due to the deletion of endogenous *PIEZO1*. *TMEM63A* variants were introduced by site-directed mutagenesis into a *TMEM63A* ires/mCherry vector. HEK-P1KO cells were transfected with wild-type (WT) and mutant constructs and electrophysiologically assayed for stretch-activated currents in the cell-attached patch clamp mode (**Fig. 1B**). As seen for the previously identified variants, mechanical activation by negative pipette pressure in the recording electrode did not induce stretch-activated currents in cells transfected with G553V (n = 10 cells) or Y559H (n = 10 cells) (**Fig. 1B**), indicating that these variants were also LOF. In contrast, the same method of stimulation elicited stretch-activated currents in cells transfected with S538L; maximal amplitudes of currents (Imax = 17.61 ± 10.83 pA, n = 8 cells) were of similar magnitude to those of the WT channel (Imax = 24.27 ± 17.54 pA, n = 18 cells) (**Fig. 1B**). Additionally, we observed slower deactivation kinetics in S538L stretch-activated currents relative to WT (143.3 ± 71.26, n= 7 vs. 77.48 ± 14.77, n= 7 cells, p = 0.05), suggesting that S538L was a gain-of-function mutation (**Fig. 1B**). Intriguingly, S538L appears to have a later-onset phenotype than the other HLD-associated *TMEM63A* variants ^7^.

**Fig. 1.**
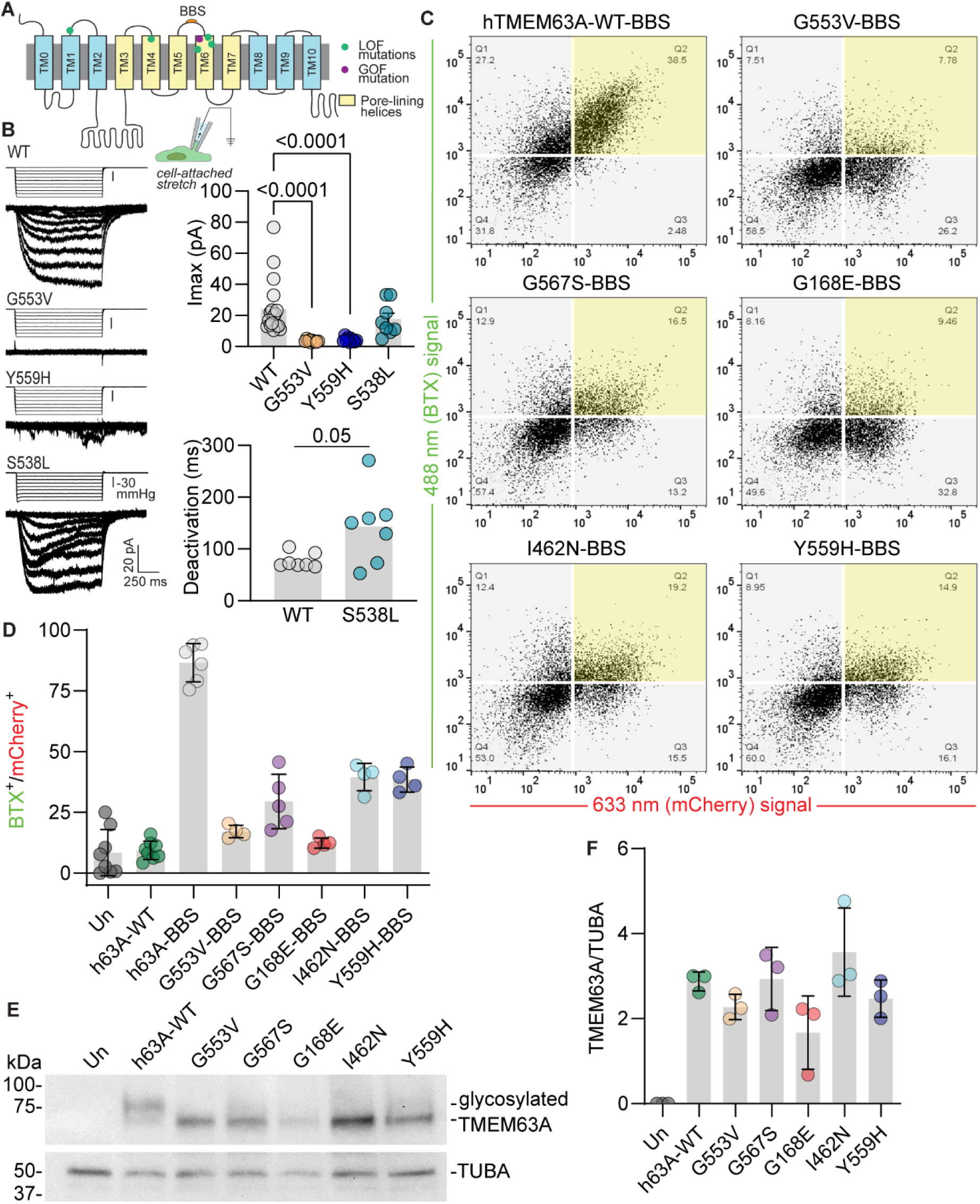
HLD19-associated *TMEM63A* LOF mutations prevent membrane localization. **(A)** Topology map of TMEM63A indicating LOF and GOF mutation sites and alpha-bungarotoxin (BTX) binding sequence (BBS). **(B)** Left, representative traces of stretch-activated currents induced by negative pipette pressure from HEK-P1KO cells transfected with WT *TMEM63A* or indicated mutant. Pressure stimulus trace is illustrated above. Right top, Maximal stretch-activated currents (I_max_) recorded from WT and variants (n = 8-18 cells per construct, p-values determined by one-way ANOVA followed by Kruskal-Wallis test of multiple comparisons). Right bottom, Quantification of deactivation kinetics measured at -80mmHg pressure stimulus (n = 7, Mann-Whitney test). **(C)** Representative flow cytometry data plots for cells transfected with WT *TMEM63A* or indicated mutant, with BBS. Transfected cells identified by mCherry fluorescence (633 nm signal). Surface TMEM63A binding identified by BTX-AF 488 fluorescence (488 nm signal). Region of overlap between BTX-labeled and transfected cells highlighted in yellow. **(D)** Percentage of gating events overlapping between BTX-labeled and transfected cells for WT (with and without BBS) and indicated mutants (with BBS) (n = 4-8 transfections per construct, p-values for all comparisons between WT-BBS and mutant-BBS is <0.0001, determined by one-way ANOVA followed by Kruskal-Wallis test of multiple comparisons). **(E)** Representative immunoblots of lysates prepared from cells transfected with WT *TMEM63A* or indicated mutant, probed against TMEM63A or TUBA (loading control). **(F)** Quantification of TMEM63A immunoblot signal density, normalized to TUBA signal density (n = 3 transfections per construct).

To test whether the absence of stretch-activated currents for the HLD-associated LOF variants was due to disrupted channel function or a deficit in channel trafficking, WT and mutant constructs were tested for surface localization. This assay was performed for the novel mutants, G553V and Y559H, in addition to those previously characterized for stretch-activated currents in Yan et al. (2019), G567S, G168E, and I463N. S538L was excluded from this assay as the observation of stretch-activated currents demonstrates that this channel variant was present at the cell surface. An alpha-bungarotoxin (BTX) binding sequence (BBS) was introduced by site-directed mutagenesis into an extracellular loop after residue 540 (**Fig. 1A**) in the WT and mutant constructs; the addition of the BBS at this residue allows labelling of channel proteins at the surface of non-permeabilized cells with fluorescent BTX conjugates ^26^. HEK-P1KO cells were transfected with WT (with and without BBS) and mutant *TMEM63A* ires/mCherry constructs and labelled with AF488-conjugated BTX before being subjected to fluorescence-activated cell sorting (FACS). FACS gates were set to exclude dead cells (DAPI-positive) before mCherry fluorescence (transfected cells) was compared to AF488 BTX fluorescence (surface-labelled cells) (**Fig. 1C, D**). All BBS-tagged mutants tested (I462N: 39.51 ± 5.67, n = 4 cultures; G168E: 12.35 ± 2.05, n = 4 cultures; G553V: 17.06 ± 2.59, n = 4 cultures; Y559H: 38.51 ± 5.14, n = 4 cultures; G567S: 29.46 ± 11.21, n = 5 cultures, p<0.0001) displayed 2.2 – 7.0-fold lower levels of surface-labelled cells relative to BBS-tagged WT (86.63 ± 7.85, n = 6 cultures, p<0.0001) (**Fig. 1D**). The percentage of surface-labelled cells for G553V and G168E was indistinguishable from that of untagged WT (9.37 ± 3.77, n = 8 cultures) and untransfected cells (8.41 ± 9.54, n = 8 cultures). The tested HLD19-associated *TMEM63A* variants thus significantly impair membrane localization.

The absence of surface-presenting mutant TMEM63A protein was further investigated by immunoblotting of the protein product in lysates of WT and mutant transfected HEK-P1KO cells (**Fig. 1F, G**). All variants expressed at similar levels to the WT protein (**Fig. 1G**), suggesting that these proteins were not subject to nonsense-mediated decay. Variant proteins ran at a notably lower apparent weight than the WT, likely explained by post-translational modification of WT TMEM63A, which is known to be glycosylated ^27^. These results indicate that hypomyelinating mutations in TMEM63A disrupts folding/trafficking of the channel leading to LOF.

### *Tmem63a* deletion in mice recapitulates the transient HLD19 hypomyelination phenotype

The potential etiology of HLD19 was explored by evaluating the effect of *Tmem63a* deletion in a mouse model. *Tmem63a* global knockout (KO) mice were revived from frozen embryos acquired from Nanjing Biomedical Research Institute via the International Mouse Phenotyping Consortium (IMPC). KO of *Tmem63a* was validated by RT-qPCR (**Extended Data Fig. 1A**). Given the clear effect of HLD19 on white matter tracts in the observed patients, we examined myelin-/OL-rich tissues and focused on a developmental window, postnatal days 10- 30 (P10-P30), likely to capture the transient nature of the hypomyelination experienced in the LOF patient cohort.

To test whether the mouse model recapitulated the patient phenotype, myelin output was measured in *Tmem63a^KO/KO^* mice. Myelin coverage in motor cortex was determined using myelin basic protein-positive (MBP+) area from P10, P14, P21, and P30 *Tmem63a^WT/WT^*and *Tmem63a^KO/KO^* mice (**Fig. 2A, B**). Myelin coverage was transiently reduced, with a 1.7-fold decrease in P14 *Tmem63a^KO/KO^*mice compared to WT coverage (KO: 0.033 ± 0.014, n = 6 animals; WT: 0.056 ± 0.018, n = 6 animals; p = 0.040), but fully recovered by P30 (KO: 0.18 ± 0.067, n = 4 animals; WT: 0.18 ± 0.072, n = 4 animals; p = 0.99) (**Fig. 2B**). Possible explanations for the shared myelin deficit in the mouse model and the HLD19 patients were then explored.

**Fig. 2.**
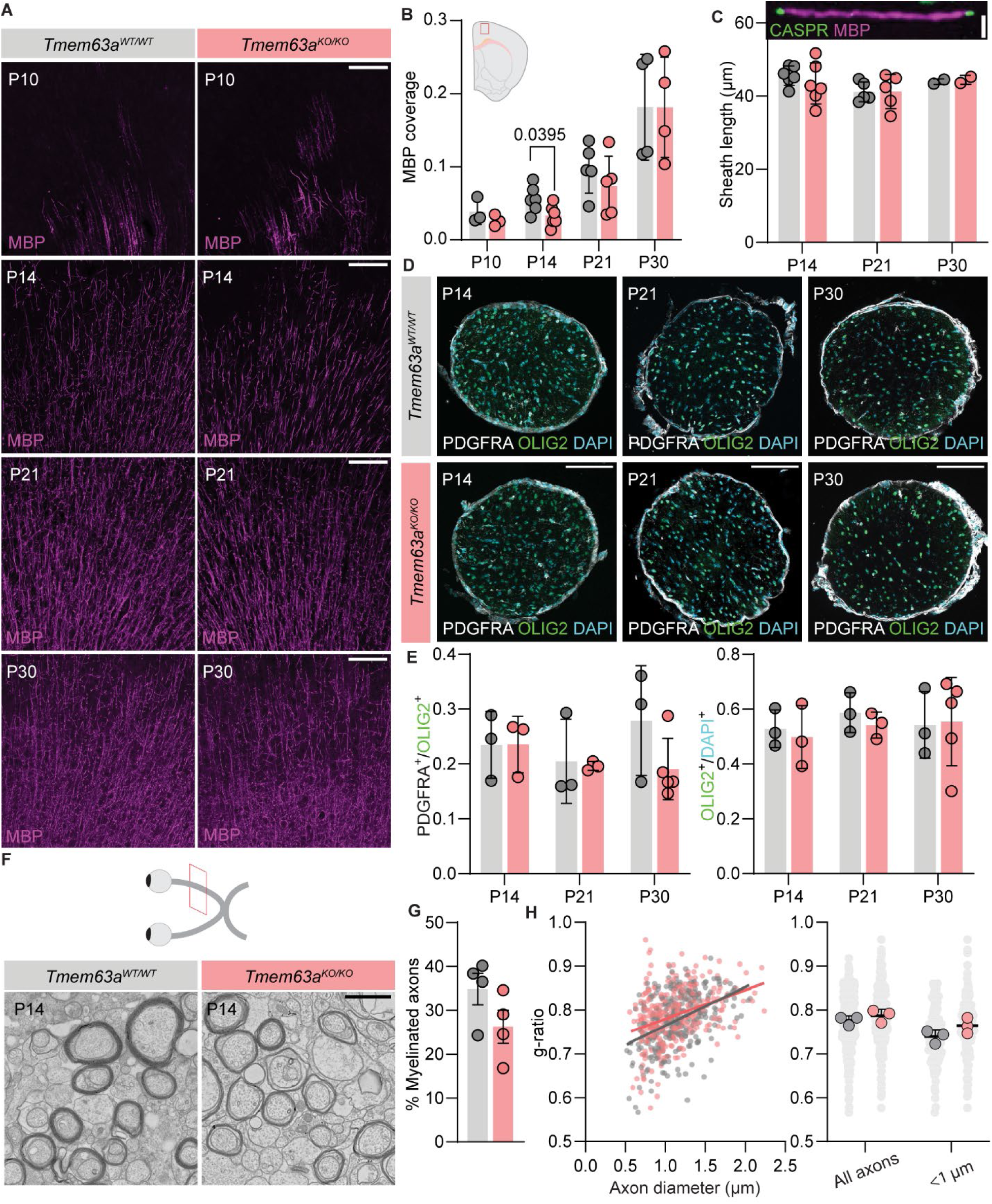
Delayed myelination in *Tmem63a*-null mice recapitulates the HLD19 phenotype. **(A)** Representative micrographs of 50 µm-thick cortical sections immunostained against MBP (magenta) from *Tmem63a^WT/WT^* (gray labels) and *Tmem63a^KO/KO^* (rose labels) mice at indicated ages. Scale bar: 100 µm. **(B)** MBP coverage in cortical sections, as MBP+ area by total area, for *Tmem63a^WT/WT^* and *Tmem63a^KO/KO^*mice. Inset region of MBP measurement in cortex (n = 4-6 animals per genotype and age, Brown-Forsythe and Welch ANOVA for multiple comparisons). **(C)** Internode myelin sheath lengths as MBP+ distance between CASPR labelling (inset), for *Tmem63a^WT/WT^* and *Tmem63a^KO/KO^* mice. (n = 4-6 animals per genotype and age, One-way ANOVA for multiple comparisons). Scale bar: 5 µm. **(D)** Representative micrographs of 14 µm-thick optic nerve sections immunostained against OLIG2 (green) and PDGFRA (gray), and counterstained with DAPI (cyan) from *Tmem63a^WT/WT^* and *Tmem63a^KO/KO^* mice. Scale bar: 100 µm. **(E)** Top, Percentage PDGFRA+ cells/OLIG2+ cells in optic nerve for *Tmem63a^WT/WT^* and *Tmem63a^KO/KO^* mice. Bottom, percentage OLIG2+ cells/DAPI+ cells in optic nerve for *Tmem63a^WT/WT^* and *Tmem63a^KO/KO^* mice (n = 3-5 animals per genotype and age, Brown-Forsythe and Welch ANOVA). **(F)** Representative transmission electron micrographs of 70 nm-thick optic nerve sections from P14 *Tmem63a^WT/WT^* and *Tmem63a^KO/KO^* mice. Scale bar: 1 µm. **(G)** Percentage of myelinated axons in P14 optic nerves from *Tmem63a^WT/WT^* and *Tmem63a^KO/KO^* mice (n = 4 animals per genotype, unpaired t-test). **(H)** g-ratio of axons from P14 optic nerve of *Tmem63a^WT/WT^* and *Tmem63a^KO/KO^*mice. Left, g-ratio by axon caliber. Right, Animal averages of g-ratios. Individual axons in light gray, animal averages in gray (*Tmem63a^WT/WT^*) and rose (*Tmem63a^KO/KO^*). (n = 3 animals per genotype, unpaired t-test).

Myelin sheath length, a feature of OL morphology, was assessed. Sheath length was determined from the same motor cortex image set used for myelin coverage by measuring internode distance along a contiguous MBP+ myelin sheath capped on both ends by axonal Contactin-associated protein (CASPR) signal (**Fig. 2C**). P10 images were omitted from this analysis due to the lack of mature myelin internodes at this timepoint. No differences in sheath length were observed between *Tmem63a^WT/WT^* and *Tmem63a^KO/KO^* mice at any age (**Fig. 2C**). To determine whether the myelin deficit might be due to a reduction in the available number of myelinating OLs, the relative proportion of OL progenitor cells (OPCs) to all OL-lineage cells (OLCs) and the relative proportion of all OLCs to total number of cells was determined in optic nerves from P14, P21, and P30 *Tmem63a^WT/WT^* and *Tmem63a^KO/KO^*mice (**Fig. 2D, E**). Optic nerve tissue was selected for this analysis due to its relatively homogeneous cell population.

OPC/OLC and OLC/total cells were determined by counting the number of platelet-derived growth factor receptor alpha (PDGFRA+; OPCs), oligodendrocyte transcription factor 2 (OLIG2+; OLCs), and DAPI+ (total cells) in immunostained optic nerve sections (**Fig. 2D**). No differences in either the OPC or OLC populations were found at any age (n = 3 – 5 animals per age, genotype) (**Fig. 2E**).

To examine subtleties in myelin morphology that may not be evident at the light-microscopy level, myelin ultrastructure in optic nerves from P14 *Tmem63a^WT/WT^* and *Tmem63a^KO/KO^* mice was evaluated by transmission electron microscopy (TEM) (**Fig. 2F-H**). TEM visualization of the optic nerve allowed determination of the percentage of myelinated axons for each genotype and the measurement of myelin g-ratio (**Fig. 2H**). The percentage of myelinated axons trended towards a decrease in P14 *Tmem63a^KO/KO^* mice (WT: 34.85 ± 7.17 %, n = 4 animals; KO: 26.27 ± 7.5 %, n = 4 animals; p = 0.15). g-ratio of all axons was unchanged (WT: 0.78 ± 0.010, n = 3 animals; KO: 0.79 ± 0.015, n = 3 animals; p = 0.36), although we observed a trend towards thinner myelination of small caliber axons, consistent with what has been reported previous for other hypomyelination phenotypes ^28–30^ (WT: 0.73 ± 0.014, n = 3 animals; KO: 0.76 ± 0.017, n = 3 animals; p = 0.14).

These data demonstrate that *Tmem63a^KO/KO^* mice recapitulate the transient hypomyelination observed in human phenotype. The mice have a clear reduction in cortical MBP+ coverage, suggesting a lag in myelin formation. The observation of fewer myelinated axons in the optic nerve further suggests that the myelin deficit in the absence of *Tmem63a* may be attributed to an initial failure of axonal selection.

### The function of TMEM63A in myelination is conserved in zebrafish

To test evolutionary conservation of TMEM63A across the vertebrate lineage, myelin was evaluated in zebrafish, the simplest laboratory model with myelin. To perturb *tmem63a* in zebrafish, sgRNAs targeting *tmem63a* were designed against restriction sites in *tmem63a*, such that sgRNA efficiency could be tested by restriction digest, and a highly efficient sgRNA was identified (**Fig. 3A** and **Extended Data Fig. 2**). This sgRNA and Cas9 protein were injected into one-cell zebrafish zygotes in the transgenic *Tg(mbp:egfp-caax)* background, and myelin membrane was visualized in injected F0 “crispant” animals. *mbp* intensity was measured in the dorsal and ventral spinal cord of 4- and 5- days post-fertilization (dpf) crispants (**Fig. 3B, C**).

**Fig. 3.**
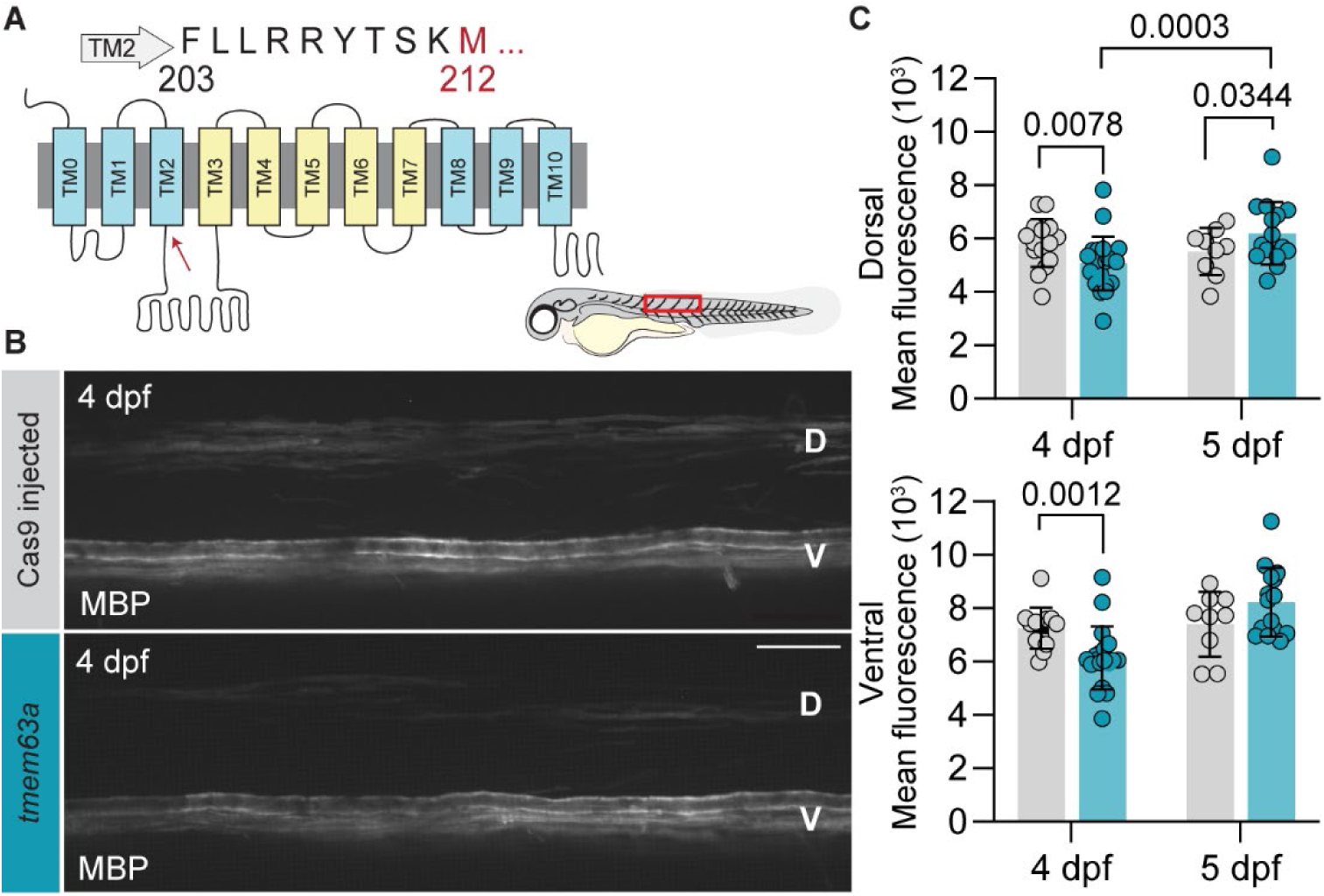
TMEM63A function in myelination is conserved in zebrafish. **(A)** Topology map of *danio rerio* Tmem63a; region targeted by CRISPR/Cas9 flanks M212 (highlighted in red and indicated by red arrow on map). **(B)** Representative micrographs comparing control Cas9-injected or *tmem63a* sgRNAs injected in *Tg(mbp:EGFP-CAAX)* fish at 4 days post-fertilization (dpf). D, dorsal and V, ventral spinal cord. Cartoon larva illustrates ROI (red box). Scale bar: 25 µm. **(C)** Mean MBP intensity of dorsal (top) and ventral (bottom) spinal cord from z-stack sum projections of Cas9 and *tmem63a* F0 fish. (n = 25 – 32 animals per crispant and age, mixed effects analysis of multiple comparison by uncorrected Fisher’s LSD).

Strikingly, as in *Tmem63a^KO/KO^* mice, *tmem63a* crispants had reduced *mbp* expression relative to control Cas9-injected fish in both dorsal (5264 AU, n = 32 *tmem63a* crispants vs 6018 AU, n = 31 *Cas9* crispants; p = 0.0078) and ventral spinal cord (6343 AU, n = 31 *tmem63a* crispants vs 7443 AU n = 30 *Cas9* crispants; p = 0.0012) at 4 dpf that recovers by 5 dpf (**Fig. 3C**). These results bolster the notion that TMEM63A is an evolutionarily conserved regulator of myelination and establish zebrafish as a powerful model to study TMEM63A in myelination *in vivo*.

### TMEM63A functions cell-autonomously in oligodendrocytes

Both bulk RNA-sequencing and single nuclei RNA-sequencing from the CNS indicate *Tmem63a* is enriched in postmitotic OLs ^31,32^. To determine whether TMEM63A influences early stages of myelination through an OL-intrinsic mechanism, *Tmem63a* knockdown exclusively in myelinating OLs was accomplished using a floxed *Tmem63a* allele (*Tmem63a^Fl^*) in conjunction with *Cre* recombinase driven by *Cnp1* (*Cnp1^Cre^*) or *Mobp* (*Mobp^iCre^*) ^33,34^. Knockdown in the conditional KO (cKO) was validated in both lines by RT-qPCR (**Extended Data Fig. 1B**).

Because the myelin deficit was only observed at P14 in the global *Tmem63a^KO/KO^* cortex, efforts analyzing cKOs were restricted to that time point. As in the global line, myelin coverage in motor cortex was determined using MBP+ area (**Fig. 4A, B**). *Tmem63a^Fl/Fl^; Cnp1^Cre/WT^* mice had 1.2-fold reduced myelin coverage relative to *Tmem63a^WT/WT^; Cnp1^Cre/WT^* mice (cKO: 0.054 ± 0.008, n = 6 animals, WT: 0.067 ± 0.007, n = 5 animals, p = 0.024), whereas sheath length was unchanged between genotypes (**Fig. 4B**). The same analyses were performed for *Tmem63a^Fl/Fl^; Mobp^iCre^* mice and *Tmem63a^Fl/Fl^; Mobp^WT^* mice at P14 (**Extended Data Fig. 1C, Extended Data Fig. 3**). There was a trend towards reduced myelin coverage in *Tmem63a^Fl/Fl^; Mobp^iCre^*mice, by 1.7-fold (cKO: 0.066 ± 0.005, n = 2 animals WT: 0.11 ± 0.028, n = 3 animals; p = 0.11) (**Extended Data Fig. 3B**).

**Fig. 4.**
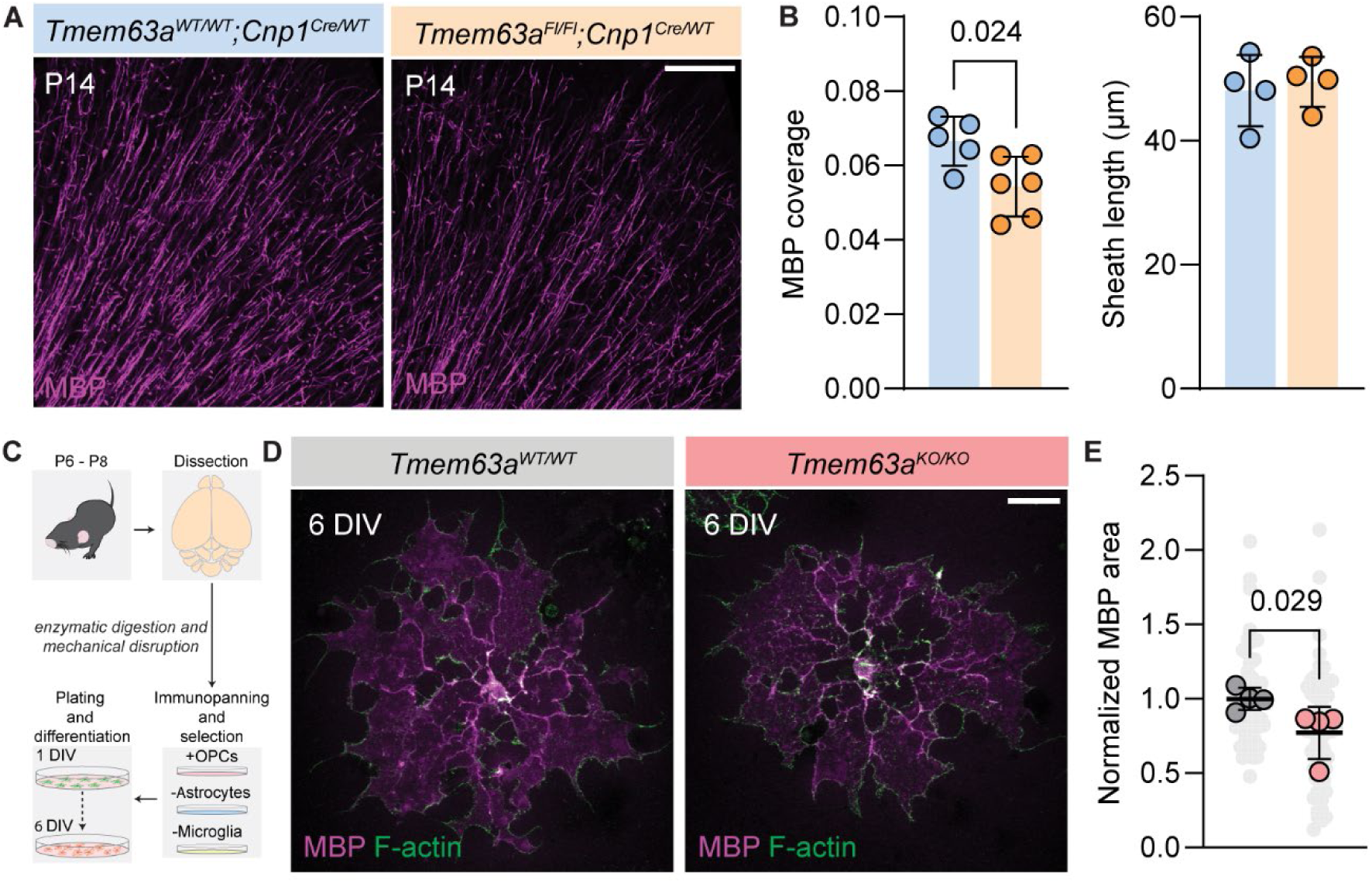
TMEM63A functions cell autonomously in oligodendrocytes. **(A)** Representative micrographs of 50 µm-thick cortical sections immunostained against MBP (magenta) from P14 *Tmem63a^WT/WT^; Cnp1^Cre/WT^* and *Tmem63a^Fl/Fl^; Cnp1^Cre/WT^*mice. Scale bar: 100 µm. **(B)** Left, MBP coverage in cortical sections, as MBP+ area by total imaged area, for *P14 Tmem63a^WT/WT^; Cnp1^Cre/WT^* and *Tmem63a^Fl/Fl^; Cnp1^Cre/WT^* mice. (n = 5-6 animals per genotype, unpaired t-test). Right, Internode myelin sheath lengths measured as MBP+ distance between nodes identified by CASPR labelling, determined for *P14 Tmem63a^WT/WT^; Cnp1^Cre/WT^* (blue) and *Tmem63a^Fl/Fl^; Cnp1^Cre/WT^* (orange) mice. (n = 4 animals per genotype, unpaired t-test unpaired t-test, 0.74). **(C)** Schematic for OL isolation and culture. After plating as OPCs, cells were differentiated for 6 days *in vitro* (DIV) before immunostaining. **(D)** Representative micrograph of OLs isolated from *Tmem63a^WT/WT^* and *Tmem63a^KO/KO^*mice, stained with MBP (magenta) and phalloidin for F-actin (green). Scale bar: 25 µm. **(E)** MBP+ area, normalized to average of WT cultures. Individual cells in light gray, culture averages in gray (*Tmem63a^WT/WT^*) and rose (*Tmem63a^KO/KO^*) (n = 4 cultures per genotype, unpaired t-test, 0.029).

Differences in OL morphology between *Tmem63a^WT/WT^* and *Tmem63a^KO/KO^* cells were next considered in OL monocultures to determine whether TMEM63A-mediated reduction in myelin was cell autonomous (**Fig. 4C-E**) ^35–37^. Consistent with observations in intact animals, *Tmem63a^KO/KO^* OLs had reduced area of MBP+ sheaths relative to *Tmem63a^WT/WT^* cells (KO cultures: 0.77 ± 0.18, n = 4; WT cultures:1.0 ± 0.07, n = 4; p = 0.029) (**Fig. 4E**).

The observed reduction in myelin coverage at P14 in both *Tmem63a* cKO mouse lines tested phenocopies the global KO, and isolated *Tmem63a^KO/KO^* OLs have impaired MBP+ output, strongly supporting an OL-autonomous role for TMEM63A in myelination.

### Myelin-associated proteins are reduced at the plasma membrane in the absence of TMEM63A

While our electrophysiology data suggest that TMEM63A exerts its influence on myelin through a role at the plasma membrane, recent publications indicate a lysosomal function for TMEM63 proteins ^22,38^. Considering this, the subcellular distribution of TMEM63A was investigated. Brains from P30 - P90 *Tmem63a^WT/WT^* and *Tmem63a^KO/KO^*mice were subjected to density-based subcellular fractionation ^39^ (**Fig. 5**). Total membranes, inclusive of lysosomes, were isolated; from this fraction, endoplasmic reticulum and plasma membranes (ER/PM) were further purified. Subcellular fractions were subjected to analysis by immunoblotting and probed for the lysosomal/microsomal marker lysosome-associated membrane protein 2 (LAMP2), MBP, and TMEM63A (**Fig. 5A-D**). LAMP2 was detected across all fractions analyzed, with the lowest observed level in the purified ER/PM fraction; its presence in the cytosolic fraction likely represents a contribution from low density microsomes ^40^ (**Fig. 5B**). MBP and TMEM63A were enriched in the purified ER/PM fraction (**Fig. 5C, D**), whereas TMEM63A protein was not detected in any fraction from *Tmem63a^KO/KO^*animals (**Fig. 5D**). Taken together, the enrichment of TMEM63A in the ER/PM fraction and overlap of TMEM63A and LAMP2 across subcellular compartments indicate the existence of both plasma membrane and lysosomal populations of TMEM63A.

**Fig. 5.**
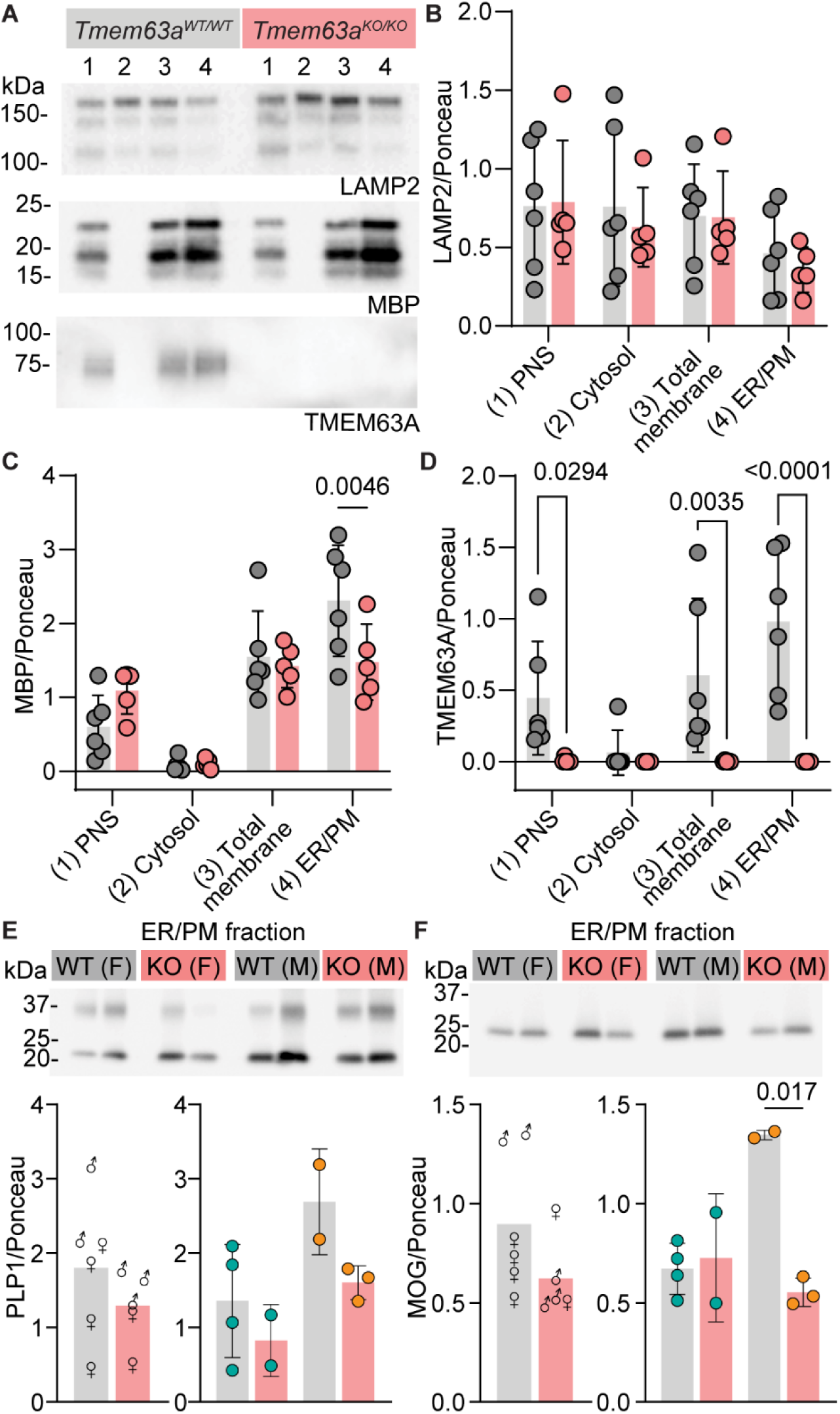
TMEM63A targets to membrane and modulates membrane-associated myelin proteins. **(A)** Representative immunoblots of fractions isolated from brain tissue of P30 – P90 *Tmem63a^WT/WT^* and *Tmem63a^KO/KO^*mice. Blots probed for lysosomal protein LAMP2, myelin-associated protein MBP, and TMEM63A. Lanes: (1) post-nuclear supernatant [PNS], (2) cytosol, (3) total cellular membranes [inclusive of ER/PM and lysosomes], (4) ER/PM. Quantification of LAMP2 **(B)** MBP **(C)** TMEM63A **(D)** immunoblot signal density normalized to Ponceau S signal density measured between 100 – 150 kDa.(n = 5-6. 2-way ANOVA and Tukey’s multiple comparison testing). **(E-F)** Top, ER/PM fractions isolated from P30 – P90 *Tmem63a^WT/WT^* and *Tmem63a^KO/KO^* mice, example immunoblot probed against PLP1 (E) and MOG (F). Bottom, Normalized immunoblot density for PLP1 (E) and MOG (F) for all animals and separated by male (orange) and female (teal) (n = 5-6 animals, one-way ANOVA).

Notably, ER/PM MBP levels were significantly lower by 1.6-fold in fractions from *Tmem63a^KO/KO^* mice than those from *Tmem63a^WT/WT^*mice (KO:1.48 ± 0.51, n = 5 animals, WT:2.31 ± 0.75, n = 6 animals; p = 0.0046). This led us to investigate the levels of other myelin-membrane associated proteins, proteolipid protein 1 (PLP1) and myelin oligodendrocyte glycoprotein (MOG) in the purified ER/PM fraction of these animals (**Fig. 5E, F**). By immunoblotting, PLP1 and MOG trend towards reduced levels in *Tmem63a^KO/KO^* mice.

Curiously, differences are more exaggerated for PLP1 and MOG in ER/PM fractions from male animals (**Fig. 5E, F**). The data suggest that TMEM63A regulates myelin production by OLs, which presents as reduced myelin coverage in the absence of *Tmem63a* or inactive variants of *TMEM63A.* These results also imply that TMEM63A could modulate myelin production through downstream effects on major myelin protein constituents.

### Surface expression of TMEM63A is short-lived

The existence of both plasma membrane and lysosomal populations of TMEM63A raised the question if TMEM63A proteins might be cycling between these cellular compartments. Thus, the longevity of TMEM63A at the extracellular membrane was interrogated. Returning to the BBS-engineered *TMEM63A* constructs used to first investigate the protein’s localization on cell surfaces, WT BBS-tagged TMEM63A was assayed for internalization (**Fig. 6A, B**). Live HeLa P1KO cells were labelled with AF-647-conjugated BTX and either immediately fixed (0 h) or returned to 37°C for 2 h incubation and then fixed (2 h) (**Fig. 6A**). Abundant surface labelling of BBS-tagged TMEM63A (*h63a(WT)*-*BBS*) with BTX was observed at 0 h (+BTX cultures:91.05 ± 35.25 AU, n = 3 0h, -BTX cultures: 19.02 ± 24.33 AU, n = 3 0h; p = 0.029), but BTX signal disappeared after 2 h (+BTX cultures:28.85 ± 21.16 AU, n = 3 2h, -BTX cultures:18.11 ± 11.27 AU, n = 3 2h; p = 0.95). On the contrary, as expected the LOF variant *TMEM63A(I462N)* showed no surface labeling at either time point (**Fig. 6C, D**). Because internalized fluorescent BTX conjugates have previously been reported ^41^, the absence of BTX signal at this time point could suggest that the conjugate’s fluorescence was quenched by an acidic environment such as the lysosomal compartment. In support of this interpretation, the internalization assay was repeated for BBS-tagged mouse PIEZO2 (*Piezo2-BBS*, inserted after amino acid 568) in HEK- P1KO cells, which were more amenable to *Piezo2* expression than HeLa-P1KO cells (**Fig. 6E, F**). While BTX labelling of PIEZO2-expressing cells appeared sparser than BTX labelling of TMEM63A-transfected cells, the BTX labelling remained elevated at 2 h (+BTX cultures:75.65 ± 23.46 AU, n = 3 2h, -BTX cultures:19.77 ± 14.25 AU, n = 3 2h; p = 0.18), although it decreased from 0 h levels (+BTX cultures:114.5 ± 52.65 AU, n = 3 0h, -BTX cultures: 8.67 ± 4.79 AU, n = 3 0h; p = 0.010) (**Fig. 6F**). These data suggest that TMEM63A proteins might rapidly cycle through phases at the plasma membrane and inside lysosomes/lysosomal membranes. TMEM63A may have discrete roles on cell surfaces and in lysosomes.

**Fig. 6.**
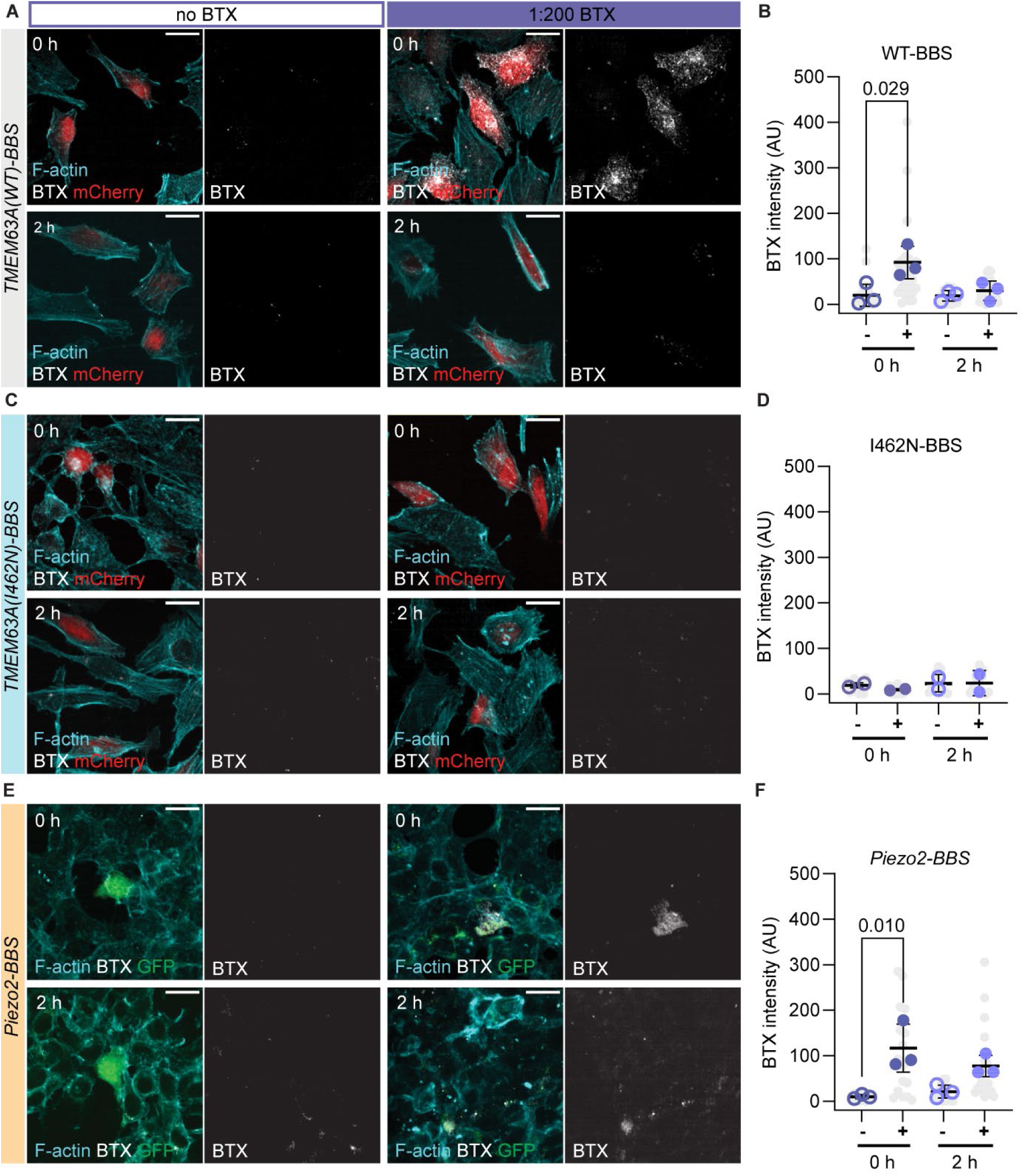
Surface-localized TMEM63A internalizes rapidly. **(A)** Representative micrographs of HeLa-P1KO cells transfected with *TMEM63A(WT)-BBS/ires/mCherry* prior to incubation with BTX-AF 647. Cells were fixed either immediately after (0h) or 2h after removal of BTX conjugate (2h). BTX signal (grays), F-actin (cyan), and mCherry (red) are shown. Scale bar: 20 µm. **(B)** Quantification of BTX signal at 0 and 2h post labelling for conditions with (solid markers) or without (empty markers) conjugate for *TMEM63A(WT)-BBS*. Individual cells shown in gray, culture/transfection averages shown in blue (n = 3 transfections, one-way ANOVA by Tukey’s multiple comparison). **(C)** Representative micrographs of HeLa-P1KO cells transfected with *TMEM63A(I462N)-BBS/ires/mCherry* prior to incubation with BTX-AF 647 conjugate, as in (A). **(D)** Quantification of BTX signal at 0 and 2h post labelling for conditions with (solid markers) or without (empty markers) conjugate, as in (B) (n = 2 transfections). **(E)** Representative micrographs of HEK-P1KO cells transfected with *Piezo2-BBS/ires/EGFP* prior to incubation with BTX-AF 647 conjugate, as in (A). **(F)** Quantification of BTX signal at 0 and 2h post labelling for conditions with (solid markers) or without (empty markers), as in (B) (n = 3 transfections, one-way ANOVA by Tukey’s multiple comparison).

## DISCUSSION

The data discussed here strongly support that TMEM63A regulates myelin formation. The loss of *Tmem63a* results in reduced myelination in global mouse mutants and zebrafish crispants, suggesting a conserved role across species. In mice, isolated OLs and OL-specific knockdown of the channel reproduce phenotypes observed in global mutants and human patients, pointing to a defined OL-role for TMEM63A. Furthermore, TMEM63A is present on plasma membrane and lysosomes, and co-segregates to these compartments with OL-expressed, myelin-associated proteins, defining its cellular localization in OLs.

The LOF HLD19-associated *TMEM63A* variants have impaired surface targeting (**Fig. 1** and **Fig. 6**). This supports the premise that TMEM63A modulates myelination via activity at OL surfaces because the decrease in the number of functional channels at cell surfaces in the HLD19 cohort is sufficient to induce hypomyelination. Indeed, the studies of the mouse *Tmem63a* KO indicate that a complete absence of TMEM63A also induces transient hypomyelination.

Intriguingly, while the myelin deficit observed in the mouse model appears to resolve rapidly, adult (∼P60) mice lacking *Tmem63a* exhibit abnormal gait (IMPC). The development of abnormal gait in older animals suggests development of new/further myelin impairments after the resolution of the original deficit observed at P14, as motor difficulties are frequently a symptom of myelin dysfunction ^42,43^. The HLD19 patient cohort may reflect this potential recurrence of a myelin deficit. The subject described by Garg et al. (2024) experienced sudden onset of symptoms at age 31; while no history of neurological symptoms was described, this patient was not assessed at the same young ages as the other members of the cohort, leaving it possible that she may have also experienced a clinically missed period of mild hypomyelination. Conversely, the younger patients evaluated for leukodystrophy may present further symptoms as adults. The difference in onset ages could also reflect an inherent difference in the consequences of the GOF mutation discovered by Garg et al., which retains mechanosensitive channel activity, and the LOF mutations found in the other patients.

The experiments in mice and zebrafish, along with data from the HLD19 cohort, clearly demonstrate that TMEM63A modulates myelin formation (**Fig. 2, 3**) and as a mechanosensor, TMEM63A activity on OL surface membranes might be responsible for the channel’s effect on myelin ^44^. This raises the important question of what specific mechanical cues activate TMEM63A on OLs. During axonal selection and the ensuing generation of myelin wraps, OL processes must be recognizing specific amount of membrane stretch and corresponding changes in membrane tension. A prevailing hypothesis is that a mechanosensitive channel(s) that is permeable to Ca^2+^, like TMEM63A, transduces these membrane changes ^45,46^. In zebrafish, OL process extension and retraction are accompanied by distinct Ca^2+^ signals ^47,48^, supporting a model where these events are mediated by mechanosensitive channels with unique biophysical properties that can regulate the amount of Ca^2+^ for each event and in turn govern subsequent signaling in OL morphology and function. Therefore, one can hypothesize that TMEM63A- mediated Ca^2+^ influx in response to membrane stretch during contact formation and myelin extension cues the synthesis of more myelin and elongation of the process. If true, axonal selection for myelination may be the bottleneck encountered in hypomyelination due to the lack of functional TMEM63A channels. This is supported by the trend towards fewer myelinated axons in the P14 knockout optic nerve (**Fig. 2H**). Together, our data conclusively demonstrate that TMEM63A is well-positioned to sense stretch during active myelination and regulate myelin formation.

In addition to localization at the plasma membrane, our data also suggest the presence of TMEM63A in membranous intracellular compartments, inclusive of lysosomes (**Fig. 5**). A lysosomal function of TMEM63A would further explain the hypomyelinating phenotype, as well as be consistent with previous reports of TMEM63A’s biological activity ^22,38,49^. Lysosomal function plays an important role in myelin formation, as lysosome-mediated autophagy is a critical component of myelin compaction and maintenance ^50^. Lysosomal vesicular trafficking contributes to membrane addition and expansion during myelin sheath growth and maturation ^51^. This is exemplified by the evidence that lysosomal exocytosis enables trafficking of PLP1 and cholesterol clusters to the OL membrane, and disruption of exocytosis results in accumulation of PLP1 in intracellular vesicles ^52^. Interestingly, purified brain membrane from *Tmem63a* KO mice has lower levels of PLP1 and MOG than wildtype membranes (**Fig. 5E, F**); a lysosomal role for TMEM63A may mediate its downstream effects on other constituent myelin proteins. Unsurprisingly, many lysosomal disorders also cause hypomyelination ^53–55^.

While TMEM63A can signal in all the modalities described above, OLs also express other mechanosensitive ion channels that may complete these roles either separately or complementarily to TMEM63A. Another member of the TMEM63 family, TMEM63B, is expressed in OLs, as is PIEZO2 ^31,56^. Future work will assess the intersection of functionality between different OL mechanosensors. In conclusion, we provide novel insights into the biological processes underlying HLD19, demonstrating that the mechanosensitive ion channel TMEM63A is a key regulator of myelination. The data presented here also support a non-organellar function for this channel, with evidence that TMEM63A is active at OL surface membranes. Furthermore, multiple vertebrate models of hypomyelination were established in mouse and zebrafish. All models will prove invaluable in the advancement of the understanding of *in vivo* myelination, as well as in the development of drug therapies for leukodystrophies.

## METHODS

### Animals

*Mice:* All mouse procedures were approved by the IACUC at Oregon Health and Science University (OHSU) (protocol IP00002957). Both female and male mice were used for all experiments with every effort made to balance the number of each sex used. Mouse lines have been maintained on a C57BL/6J background.

The global *Tmem63a* knockout (*Tmem63a^KO^*) mice used here were revived (at Scripps Research) from frozen embryos acquired from Nanjing Biomedical Research Institute via the International Mouse Phenotyping Consortium (IMPC; strain name: B6/N-Tmem63atm1/Nju, strain number: T000698). The insertion of Velocigene cassette ZEN-Ub1 creates a 27,792 bp deletion in the positions 182,876,714-182,904,505 of Chromosome 1. The floxed *Tmem63a* (*Tmem63a^Fl^*) mice were acquired as ES cells from EUCOMM via IMPC (MAE-4982); loxP sites flank Exon 6 of the gene. Mice were generated at Scripps Research. *Tmem63a^KO^* and *Tmem63a^Fl^* mice were transferred to OHSU under an MTA. The *Cnp1*^Cre^ line ^33^ was provided by the Monk lab. The *Mobp^iCre^* line was created in the Bergles lab ^34^; founder animals were transferred from the Popko lab. To produce *Cnp1*-conditional *Tmem63a* knockout mice, *Tmem63a^Fl/WT^* mice were crossed to *Tmem63a^Fl/WT^; Cnp1^Cre/WT^* mice. To produce *Mobp*-conditional *Tmem63a* knockout mice, *Tmem63a^Fl/WT^*mice or *Tmem63a^Fl/Fl^* mice were crossed to *Tmem63a^FL/WT^; Mobp^iCre/WT^* mice. All genotyping was outsourced to TransnetYX, Inc. (Cordova, TN).

Genotyping primers for *Tmem63a^KO/KO^* mice: To identify null gene (KO or heterozygous): Forward primer: CTT GGC CAC AAA CCA CTA GG; Reverse primer: TCA TTC TCA GTA TTG TTT TGC C. To identify WT gene (WT or heterozygous): Forward primer: GCA AGC CTG TGC TAT TTG AGA TG; Reverse primer: GTG TCC AAC TCT TCA CAC TAA CCT C. Genotyping primers for *Tmem63a^Fl^*mice: Forward primer: GAA GTG GAC AGT CAG TTC TGA ACC; Reverse primer: CGC AGA GAC ACT CTT AAC AGG TAG.

Tissues from *Tmem63a^KO^*, *Tmem63a^Fl^*; *Cnp^Cre^*, and *Tmem63a^Fl^; Mobp^iCre^* animals at P10, P14, P21, and P30 were used for assessment of myelin coverage and sheath morphology in motor cortex, determination of OPC populations in optic nerve. Optic nerves from P14 *Tmem63a^KO^*animals were used for evaluation of myelin ultrastructure. Cortices from P6-P8 *Tmem63a^KO^* animals were used for the isolation and differentiation of OPCs (described below) for wrapping and morphological assays. Cortices from adult *Tmem63a^KO^* animals were used for subcellular fractionation experiments.

*Zebrafish crispants:* Zebrafish were maintained according to institutional ethical guidelines for research at OHSU and all experiments were approved by the IACUC (protocol IP00001148). To assess how Tmem63a affects myelination in zebrafish, mosaic mutations were introduced into zebrafish larvae in the *Tg(mbp:egfp-caax)* line using CRISPR-Cas9. sgRNAs targeting *tmem63a* were designed using the CHOP CHOP web tool (https://chopchop.cbu.uib.no/; ^57^) and synthesized using MEGAshortscript T7 Transcription kit (Thermo Fisher). sgRNAs were designed so that cutting efficiency could be assessed by restriction digest. The sgRNA CGCTACACCTCGAAAATGAAGGG was determined to have greater than 95% cutting efficiency (**Extended Data Fig. 2**) and was used for this crispant experiment. sgRNA was mixed with Cas9 Nuclease (Integrated DNA Technologies) for a final concentration of 90 ng/ml of sgRNA and 1 mg/ml of Cas9 protein. Fertilized embryos at the single-cell stage were injected with 1-2 nl of the sgRNA/Cas9 solution or Cas9 (1mg/ml) alone.

At 4- and 5 days post fertilization, transgenic fish were anesthetized with 600 mM tricaine (TRS5, Pentair) and mounted laterally using 1.2% low melting point agarose (A9414, Sigma). Fish were imaged at body segment 7 on a Zeiss spinning disc confocal microscope using a 20x dipping objective.

### Cell lines

Electrophysiology recordings, membrane targeting, and relative expression levels of WT and HLD19 mutants were assessed in HeLa-P1KO and HEK-P1KO cells, generous gifts from Drs. Patapoutian and Grandl, respectively. HEK-P1 KO cells were generated by the Duke Functional Genomics Core and genomically validated to have a 34 bp deletion (frameshift) in exon 42 of *PIEZO1.* Cells were cultured in DMEM (Gibco, cat# 11965-092) supplemented with 10% FBS (Gibco, cat# A5256801) and 100 U/ml penicillin-streptomycin (Gibco, cat# 15140148) at 37°C in 5% CO_2_. All experiments were performed between passages 4 and 19. Cultures were confirmed mycoplasma-negative by the Universal Mycoplasma Detection kit (ATCC, cat# 30- 1012K).

### Transfections

For electrophysiology recordings, HEK-P1KO cells were grown in Dulbecco’s Modified Eagle Medium (DMEM) containing 4.5 mg.ml^−1^ glucose, 10% fetal bovine serum, 50 units.ml^−1^ penicillin and 50 µg.ml^−1^ streptomycin. Cells were plated onto 12 mm round glass poly-D-lysine coated coverslips placed in 24-well plates and transfected using lipofectamine 2000 (Invitrogen) according to the manufacturer’s instruction. All plasmids were transfected at a concentration of 400 ng.ml^−1^. Cells were recorded from 24 to 48 h after transfection.

For bungarotoxin labeling experiments, HeLa-P1KO or HEK-P1KO cells were grown under identical culture and transfection conditions as described above.

### Isolation, culture, and differentiation of OPCs from neonatal mouse cortices

Primary mouse OPCs were isolated from P6-P8 mouse pups as previously described (for detailed description of protocol and reagents see ^36^). Briefly, mice were anesthetized using isoflurane and decapitated. The brain was dissected out, diced into ∼1 mm^3^ cubes, and incubated in solution containing papain under 5%CO2/95% O2 gas flow at 34 degrees for 1 h. The tissue was triturated to obtain a single cell suspension. Cells were passed over a series of immunopanning dishes (negative selection using GalC and Ran2 and positive selection using O4) to obtain a highly enriched OPC population. OPCs adherent to the final immunopanning O4 dish were trypsinized and resuspended in DMEM supplemented with SATO medium, B-27 (2%, Thermo Fisher) NT3 (1 ng/ml), CNTF (10 ng/ml), PDGF-AA (10 ng/ml) and expanded for 4-6 days. For morphological analysis, cells were seeded on PDL coated coverslips in 24 well plates at a density of 9000 cells/coverslip in the same media but omitting PDGF-AA and adding T3 (50 nM) to promote differentiation. Cells were maintained in an incubator at 10% CO_2_ and media was changed every other day.

### Electrophysiology

Patch-clamp experiments in HEK-P1KO cells were performed in standard cell-attached mode using Axopatch 200B amplifier (Axon Instruments). Currents were sampled at 20 kHz and filtered at 2 kHz or 10 kHz. Leak currents before mechanical stimulations were subtracted off- line from the current traces. All experiments were performed at room temperature (RT). To induce stretch-activated currents, membrane patches were stimulated with 1 s negative pressure pulses through the recording electrode using Clampex controlled pressure clamp HSPC-1 device (ALA-scientific), with inter-sweep duration of 1 min and subsequent sweeps applied at a delta of 10mmHg. External bath solution used to zero the membrane potential consisted of (in mM) 140 KCl, 1 MgCl_2_, 10 glucose and 10 HEPES (pH 7.3 with KOH). Recording pipettes were of 1 – 3 MΩ resistance when filled with standard solution composed of (in mM) 130 mM NaCl, 5 KCl, 1 CaCl_2_, 1 MgCl_2_, 10 TEA-Cl and 10 HEPES (pH 7.3 with NaOH).

### Flow cytometry

To assess surface expression of HLD19-associated TMEM63A mutants, fluorescence-activated cell sorting (FACS) was used to identify membrane-labelling with fluorescent α- bungarotoxin (BTX) of wildtype and mutant TMEM63A carrying a BTX-binding site introduced after residue 540. Transfection of HEK293T-P1KO cells was performed using Lipofectamine 2000 according to the manufacturer’s instructions and cells were allowed to express for 48 h. In addition to *Tmem63a* sequences, plasmids also carried an IRES element and mCherry sequence to facilitate the identification of transfected cells. Cells were resuspended in versene [PBS, 0.02% EDTA] and transferred to 1.7 mL microcentrifuge tubes. Cells were pelleted by centrifugation for 5 minutes at 200 x g, and supernatant was discarded. Cell pellets were resuspended in 100 uL per final sample of FACS buffer [PBS, 2% FBS, 1 mM EDTA] depending on staining condition either with or without Alexa Fluor 488-conjugated BTX (Invitrogen, cat# B13422) at 1:100, and transferred to a 96-well plate. Plates were protected from light with aluminum foil and incubated at RT for 15 minutes. Cells were washed 3x by centrifugation for 5 minutes at 200 x g, removal of supernatant by quick inversion, and resuspension in FACS buffer. Cells were then resuspended again in FACS buffer depending on staining condition either with or without DAPI at 1:5000 (VWR, cat# 89139-050) and transferred to FACS tubes on ice. Sample fluorescence was measured on a LSR II Flow Cytometer (BD Biosciences). Untransfected, unstained cells were used to standardize gates to exclude non-cell events. Doublets were excluded via a doublet discriminator, and dead cells were excluded by gating out the DAPI-positive population. Gates for positive and negative populations were set based on the corresponding populations in positive controls for each wavelength measured.

### Immunostaining

To examine *Tmem63a*-dependent deficits in myelination onset and progression, brains of P10, P14, P21, and P30 mice were collected from WT and KO animals. Prior to tissue collection, mice were deeply anesthetized by exposure to isoflurane, then perfused with phosphate-buffered saline (PBS) (Gibco, cat# 14190) followed by 4% paraformaldehyde (EMS, cat# 15710) in PBS. Isolated brain tissue was post-fixed by immersion in 4% paraformaldehyde in PBS for a minimum of 3 h. Post-fixed tissue was washed in PBS before sinking in 30% sucrose (Fisher, cat# BP220) in PBS. Sucrose-equilibrated tissue was embedded in O.C.T. (VWR, cat# 25608- 930) and flash-frozen in liquid nitrogen. Frozen fixed tissue was cut into 40 µm-thick sections using a Leica cryostat and collected into PBS. At RT, free-floating sections were serially dehydrated in ascending and descending ethanol steps (50%, 70%, 90%, 95%, 100%, 100%, 95%, 90%, 70%, 50%) before being blocked and permeabilized in 5% normal donkey/goat serum and 0.1% Triton X-100 in PBS (blocking solution). Labeling was performed overnight at 4°C with the following primary antibodies diluted in blocking solution: mouse IgG2a anti-CASPR 1:250 (Millipore Sigma, cat# MABN69), chicken anti-MBP 1:200 (Aves Labs, cat# MBP). Sections were washed extensively in PBS then incubated with secondary antibodies: goat anti-mouse IgG2a AF568 1:1000 (Thermo Scientific A21134), donkey anti-chicken IgY AF488 1:1000 (Thermo Scientific A78948). Sections were mounted with Fluoromount G with DAPI, and the entire depth of sections imaged using a Zeiss Apotome2 or an Andor BC43 with 20X objectives.

To evaluate potential differences in OL differentiation, optic nerves were collected from the same cohort of *Tmem63a^KO^* mice described above. As above, isolated optic nerves were post-fixed by immersion in 4% paraformaldehyde in PBS for 3 h. Post-fixed tissue was washed in PBS before sinking in 30% sucrose in PBS. Sucrose-equilibrated tissue was embedded in O.C.T. and flash-frozen in liquid nitrogen. Frozen fixed tissue was cut into 12-16 µm-thick sections using a Leica cryostat and thaw-mounted onto glass slides. Staining and imaging were performed largely as above, with omission of the serial dehydration steps. Primary antibodies used included: goat anti-PDGFRA 1:400, rabbit anti-OLIG2 (Millipore, AB9610, 1:200). Secondary antibodies used included: donkey anti-goat AF647 1:1000, donkey anti-rabbit AF488 1:1000.

To evaluate potential differences in cultured OLs from WT and KO mice, OPCs were isolated as described above. After 6 days in culture, culture media was removed, cells were washed once with PBS and fixed for 8 minutes with 4% PFA. Cells were washed 3 times with PBS and placed in blocking solution (5% donkey serum containing 0.1% Triton X-100) for 1 h at RT. Coverslips were incubated with chicken anti-MBP overnight at 4°C, washed 3 times with PBS, incubated with donkey anti-chicken IgY AF488 and phalloidin CF568 for 1 h at RT, and washed 3 times with PBS. Coverslips were mounted in Fluoromount G with DAPI and imaged using 40X oil objectives on a Zeiss LSM 980 with Airyscan. Images were acquired and analyzed blind to condition.

### Transmission electron microscopy (TEM)

To examine myelin ultrastructure, optic nerves from WT and KO mice were examined by TEM at P14, the age in which hypomyelination was observed in the cortex. Tissue processing for TEM was performed as largely described by ^31^. Tissue was isolated as described for immunostaining, but then post-fixed overnight in 2% PFA/2% glutaraldehyde in 1X PBS. After post-fixation, tissue was stored in 1.5% PFA/1.5% glutaraldehyde, 50 mM sucrose, 22.5 mM CaCl_2_•2H_2_O in 0.1 M cacodylate buffer for a minimum of 7 days and a maximum of 6 months before embedding. Optic nerves underwent further post-fixation in 2% osmium tetroxide (Electron Microscopy Sciences 19190) with 1.5% potassium ferrocyanide (Electron Microscopy Sciences 25154-20) using a Biowave Pro+ microwave (Ted Pella). Contrast was enhanced by en bloc staining with Uranyless (Electron Microscopy Sciences 22409) before dehydration in ethanol and embedding in Embed 812 (Electron Microscopy Sciences 14120). 0.4 µm sections were cut on a Leica UC7 ultramicrotome and stained with 1% Toluidine Blue (Fisher Scientific T161-25) and 2% sodium borate (Fisher Scientific S248-500) to screen for tissue viability. 70 nm sections were cut and mounted on Formvar/carbon-coated copper grids (Electron Microscopy Sciences FCF100-CU-50) and then counter-stained with UranyLess (Electron Microscopy Sciences 22409), followed by 3% lead citrate (Electron Microscopy Sciences 22410). Grids were imaged at 2900x on a FEI Tecnai T12 transmission electron microscope with a 16 Mpx camera (Advanced Microscopy Techniques Corp).

### Total protein extraction

To assess protein stability of human hypomyelinating leukodystrophy (HLD)-associated TMEM63A mutants, total protein was extracted from HEK293T-P1KO cells transfected with plasmids carrying wild-type or mutant *TMEM63A*. Mutations included: G168E ^1^, G553V ^4^, G567S ^1^, I462N ^1^, Y559H ^2^. Transfection was performed using Lipofectamine 2000 (Invitrogen, cat# 11668019) according to the manufacturer’s instructions and cells were allowed to express for 48 h. Transfected cells were lysed in RIPA buffer (Sigma-Aldrich, cat# 20-188). Cell debris was sedimented by centrifugation for 10 minutes at 16,000 x g, and the total protein content of the supernatant determined by BCA assay (Pierce, cat# PI23227). Aliquots of each lysate were treated with LDS (Invitrogen, cat# NP0007) prior to freezing or use.

### Subcellular fractionation of brain tissue

To determine the subcellular localization of TMEM63A, brain cortices were fractionated into constituent compartments by means of density centrifugation. Methods were modified from ^39^. Adult *Tmem63a^WT/WT^* and *Tmem63a^KO/KO^* mice (1-3 months of age) were starved for 18 h prior to euthanasia by CO_2_ asphyxiation and decapitation. Brain tissue excluding olfactory bulbs and cerebellum/brain stem was collected into ice-cold PBS (Gibco, cat# 14190) before mincing with a clean razor blade. 100 mg of minceate was then suspended in 1 mL ice-cold 0.3 M sucrose (Fisher, cat# BP220) in H_2_O supplemented with protease inhibitor cocktail (VWR, cat# M250) at 1:100 (1:10 w/v suspension). Minceate suspension was incubated on ice for 10 minutes to induce cell swelling before the addition of 100X buffer concentrate [1 M HEPES (Fisher, cat# BP310), 0.1 M EDTA (Fisher, cat# PRH5032) pH 7.3] for a final composition of ∼0.3 M sucrose, 10 mM HEPES, 1 mM EDTA pH 7.3. Suspensions were subjected to dounce homogenization with a tight-fitting, Teflon-coated pestle, targeting about 10-12 strokes per sample. After douncing, homogenates were further diluted with homogenization buffer [HB: 0.3 M sucrose, 10 mM HEPES, 1 mM EDTA pH 7.3] before spinning for 10 minutes at 500 x g to produce a post-nuclear supernatant (PNS) by sedimenting nuclei and large cellular debris. 880 µL PNS was layered on top of a two-step, discontinuous Nycodenz (Sigma-Aldrich, cat# D2158) gradient (1 mL 9.5% /440 µL 22.5%) where w/v Nycodenz solutions were prepared in HB. The PNS- Nycodenz gradient was centrifuged for 1 h at 141,000 x g at 4°C (46,000 rpm, TLS-55 rotor, Beckman Coulter Optima TLX Ultracentrifuge). After centrifugation, the uppermost portion of the gradient was taken as the cytosolic fraction (F1), while the top interface band was taken as the total membrane fraction (F2) and understood to include plasma membrane, endoplasmic reticulum, and lysosomes. F2 was diluted 1:1 with HB and layered on top of a discontinuous gradient of 1.32 mL 33% Percoll (Sigma-Aldrich, cat# P4937) and 440 µL 22.5% Nycodenz. This gradient was centrifuged for 30 minutes at 72,000 x g at 4°C (33,000 rpm, TLS-55 rotor, Beckman Coulter Optima TLX Ultracentrifuge). After centrifugation, the gradient was aspirated in 200 µL fractions (F2.1-F2.10), leaving behind the bottom 440 µL. Plasma membrane and endoplasmic reticulum banded at the upper interface (F2.6). Per Stromhaug et al. (1998) ^39^, a lysosomal fraction was expected at a lower interface (∼F2.7-F2.8); however, this second interface was only sporadically observed for fractions isolated from *Tmem63a^WT/WT^* animals. Aliquots of each fraction collected (PNS, F1, F2, F2.1-F2.10) were reserved for analysis by immunoblotting and treated with LDS (Invitrogen, cat# NP0007) prior to freezing or use.

### Immunoblotting

LDS-treated samples were thawed and boiled at 95°C for 5 minutes before loading into 1.0 mm-thick Bis-Tris 4-12% polyacrylamide gels (Invitrogen, cat# NP0323) and subjected to electrophoresis. For subcellular fractionation experiments, samples from *Tmem63a^WT/WT^*and *Tmem63a^KO/KO^* mice were run concurrently and a subset of fractions from the second ultracentrifugation step was analyzed, F2.5-F2.8, as these fractions covered any observable interface bands. Samples were loaded by equal volume (10 µL) to assess relative enrichment in each fraction of the proteins probed. For experiments with HLD mutants, 10 µg protein was loaded per lane. Typical run settings were 50 minutes at constant 175 V using 1X MOPS running buffer [50 mM MOPS (VWR, cat# 0670), 50 mM Tris (Fisher, cat# BP152), 0.1% SDS (Thermo, cat# 15525-017), 1.025 mM EDTA]. Protein from electrophoresed gels was transferred to PVDF membranes with 0.45 µM pore size (Millipore, cat# IPFL00010) by wet transfer in Towbin’s buffer [25 mM Tris, 192 mM glycine (Fisher, cat# BP381) pH 8.3, 10% methanol (Fisher, cat# A452)] for 75 minutes at 0.15 A. Transferred blots were blocked in 2.5% milk powder (RPI, cat# M17200) in TBSt [20 mM Tris pH 7.6, 150 mM NaCl (Fisher, cat# S271), 0.1% Tween-20 (Bio-Rad, cat# 1706531] for a minimum of 30 minutes and then incubated overnight in primary antibodies diluted in the milk-TBSt blocking solution. Primary antibodies for subcellular fractionation included: chicken anti-MBP (Aves Labs, cat# MBP) at 1:10,000, rabbit anti-TMEM63A (Sigma-Aldrich, cat# HPA068918) at 1:1000, mouse anti-LAMP2 (SCBT, cat# sc-18822) at 1:500. Incubation for LAMP2 and TMEM63A was performed at R.T., while incubation for MBP was performed at 4°C. Primary antibodies for experiments with HLD mutants included: rabbit anti-TMEM63A (Sigma-Aldrich, cat# HPA066936) at 1:1000, mouse anti-TUBA (Sigma-Aldrich, cat# T8203). After incubation with primary antibodies, blots were washed 3 x 5 minutes in TBSt. Washed blots were incubated with secondary antibodies between 2-3 hours at R.T. Secondary antibodies included: goat anti-chicken HRP (Fisher, cat# A16054) at 1:5000, goat anti-rabbit HRP (Fisher, cat# 31460) at 1:3000, goat anti-mouse (Fisher, cat# 31430) at 1:5000. After incubation with secondary antibodies, blots were washed 3 x 5 minutes in TBSt. Signal was developed with application of SuperSignal West Dura ECL substrate (Fisher, cat# PIA34075) (1 mL per blot) and visualized using a ChemiDoc XRS+ Imaging System (Bio-rad). Consistent exposure times were acquired for each sample set and probe: MBP signal was captured at 5 seconds, TMEM63A at 2 minutes, LAMP2 at 2 seconds. For immunoblots examining HLD mutants, TMEM63A and TUBA were probed concurrently. After visualization of ECL signal, blots were rinsed once in TBSt and then stained with Ponceau S (Sigma-Aldrich, cat# P7170) to visualize total protein load. Ponceau S-stained blots were also imaged.

## Data analysis

### Electrophysiology

All electrophysiology data was analyzed in Clampfit 11.2.

### OL population assessment

To evaluate populations of OPCs and OLCs, PDGFRA+ (OPCs), OLIG2+ (OLCs), DAPI+ (total cells) were counted in optic nerve sections. Images were acquired as z-stacks, and sum-projected before using the Cell Counter plugin in Fiji to track numbers of PDGFRA+, OLIG2+, and DAPI+ cells. OPC proportion of all OLCs (PDGFRA+/OLIG2+) and OLC proportion of all cells (OLIG2+/DAPI+) were determined. 2-5 sections per animal were examined, and final proportions were averaged across sections from each animal. Analysis was performed blind to genotype.

#### Myelin coverage in motor cortex and of isolated Ols

To assess changes in gross myelination in tissue, the extent of myelin coverage in cortical sections was measured by approximating the area of MBP coverage. In Fiji, images were thresholded and then binarized; the same threshold was applied across a set of images acquired from the same immunostaining/imaging session. Intensity of white (MBP+) pixels was measured across each binarized stack and averaged. Average MBP+ values were divided by the intensity of an equivalent area of all white pixels to determine the proportion of MBP+ pixels as a proxy for myelin coverage throughout the area analyzed. 2-4 sections per animal were examined. The same analysis was performed for individual cells in isolated OL cultures; for these experiments, knockout animals were normalized to littermate controls processed together during each independent cell isolation to account for variability between staining rounds. Analysis was performed blind to genotype for both cortex and culture experiments.

#### Myelin internode (sheath) length

Sheath length was measured in three dimensions using the Simple Neurite Tracer (SNT) plugin in Fiji. At least 40 sheaths were measured per section analyzed, and 2-4 sections per animal were examined. Analysis was performed blind to genotype.

#### TEM analysis

Percent myelinated axon in optic nerve was determined by marking all myelinated and unmyelinated axons with the Cell Counter plugin in Fiji in 10 random images from each nerve and then calculating the proportion of total axons. Imaging and analysis were performed while blinded to genotype.

#### Zebrafish

Spinal cord fluorescence was analyzed using ImageJ. Z-stacks were aligned using a custom ImageJ macro and a sum projection of the aligned spinal cord z-stack was performed. An ROI corresponding to dorsal or ventral spinal cord was positioned and the mean fluorescence intensity for the ROI was measured.

#### Immunoblot densitometry

Signal density of immunoblots was measured using the Gel Analyzer plugin in Fiji. Signals were normalized to either total protein content on blot (Ponceau S staining) or to TUBA signal.

#### Surface labeling and internalization of bungarotoxin-labelled TMEM63A

Taking the flat geometry of HeLa cells into account, surface signal was roughly approximated using total projected bungarotoxin (BTX) signal. Area and integrated density were acquired in Fiji for individual cells. Integrated density was normalized by dividing by area, and signals were background corrected by the subtraction of area-normalized integrated density acquired for a region within the field of view with no cells. Cells were selected based on presence of mCherry or EGFP to indicate transfection and were measured blind to BTX signal. 5 – 10 cells were analyzed per culture, and 2 – 3 cultures were analyzed per experimental condition.

#### Statistical analysis

Prism software (Graphpad) was used to perform all analyses.

## List of Supplementary Materials

Materials and Methods

Extended Data Fig. 1-3

## Acknowledgments

We thank Max Shryer, Dr. Daniel Orlin, and Dr. Gregory Duncan for technical assistance. Drs. Ardem Patapoutian, Dwight Bergles, and Brian Popko for mouse lines and Dr. Jorg Grandl and the Duke Functional Genomics Core for cell lines. We thank Dr. Marc Freeman for feedback on the manuscript. We are indebted to Adriana Reyes, Tia Perry, Emma Brennan, and Suhail Akram for animal care. We thank the OHSU glia community for insightful feedback on the project.

## Author contributions

Conceptualization: SEM, BE, KRM

Methodology: SEM, BE, KRM, JH, AJS

Investigation: JH, AJS, SB, SEM, TM, DS, RAD, AMC, BN, KE

Visualization: JH, AJS, SEM

Funding acquisition: SEM, BE, KRM

Project administration: SEM

Supervision: SEM, BE, KRM

Writing – original draft: JH, AJS, SB, SEM

Writing – review & editing: JH, SEM, BE, KRM, AJS, SB, TM, DS, RAD, AMC, BN, KE

## Competing interests

Authors declare that they have no competing interests.

### Data and materials availability

All data are available in the main text or the supplementary materials. Further information and requests for resources and reagents should be directed to and will be fulfilled by the Lead Contact, Swetha Murthy.

