## Supplemental methods and figures for "TMEM63A, associated with hypomyelinating leukodystrophies, is an evolutionarily conserved regulator of myelination"

**Supplementary Materials**

Materials and Methods

*RT-qPCR to validate Tmem63a knockdown in mouse lines*: RNA was isolated from brain or optic nerve tissues collected from adult mice (P30-P90) using a Quick-RNA Purification Kit (Zymogen R1057). Brain tissue was used as starting material for the global line ($Tmem63a^{KO}$), while optic nerve was used for the conditional lines with *Cnp1* and *Mobp*-driven Cre recombinase due to its higher proportion of oligodendrocytes. cDNA was synthesized using the High-Capacity cDNA Reverse Transcription Kit (Thermo 4368814) with equal ng amounts of starting RNA. Synthesized cDNA was then used to perform qPCR using predesigned Taqman gene expression assays (Thermo: mmTmem63a-FAM, 4448892_Mm00522653_m1; mmGapdh-VIC, 4331182_Mm99999915_g1) using a CFX Connect Real-time system (Bio-rad). *Tmem63a* transcript levels were analyzed alongside *Gapdh* transcript levels as a loading control, and results for gene expression were computed using the built-in software from Bio-rad. All results are shown as a percentage of the average control (WT) level of *Tmem63a* expression.

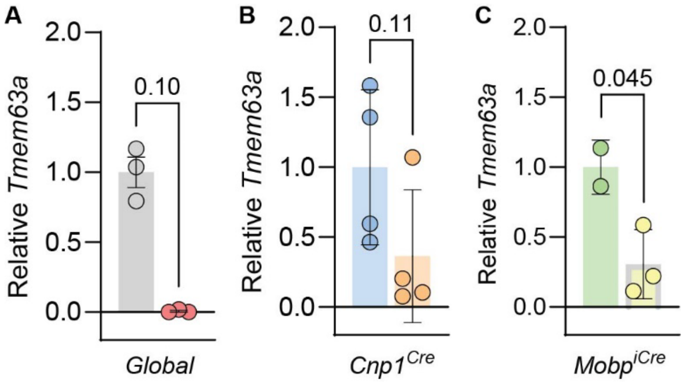


**Extended Data Fig. 1. Mouse line validation by qPCR. (A)** *Tmem63a* expression in brain tissue from adult *Tmem63a*$^{WT/WT}$ (gray, n = 3) and *Tmem63a*$^{KO/KO}$ (rose, n = 3) mice. P = 0.077. **(B)** *Tmem63a* expression in optic nerves from adult *Tmem63a*$^{WT/WT}$; *Cnp1*$^{Cre/WT}$ (blue, n = 4) and *Tmem63a*$^{Fl/Fl}$; *Cnp1*$^{Cre/WT}$ (orange, n = 4) mice. P = 0.11. **(C)** *Tmem63a* expression in optic nerves from adult *Tmem63a*$^{Fl/Fl}$; *Mobp*$^{WT}$ (green, n = 2) and *Tmem63a*$^{Fl/Fl}$; *Mobp*$^{iCre}$ (yellow, n = 3) mice. P = 0.045. (A)-(Cp-value by Mann-Whitney test).

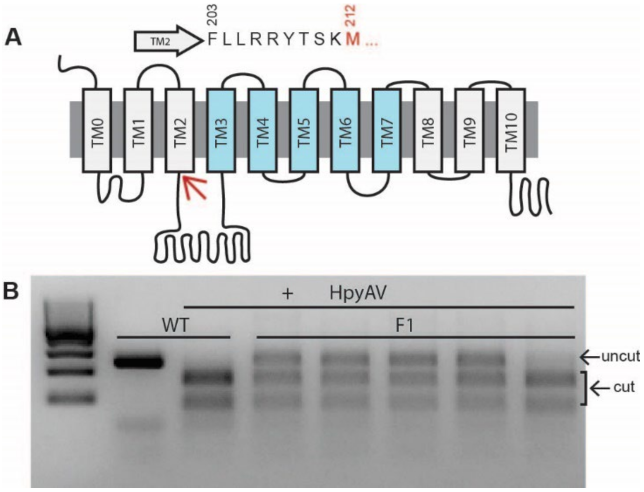


**Extended Data Fig. 2. Generation of *tmem63a* global knockout fish. (A)** Cartoon representation of *danio rerio* tmem63a with putative pore lining transmembrane domains in blue. The residue targeted through CRISPR/Cas9 is highlighted in red and indicated by the red arrow. **(B)** Agarose gel comparing wild type and the F1 generation that is heterozygous for a mutation resulting in destruction of the HpyAV restriction site.

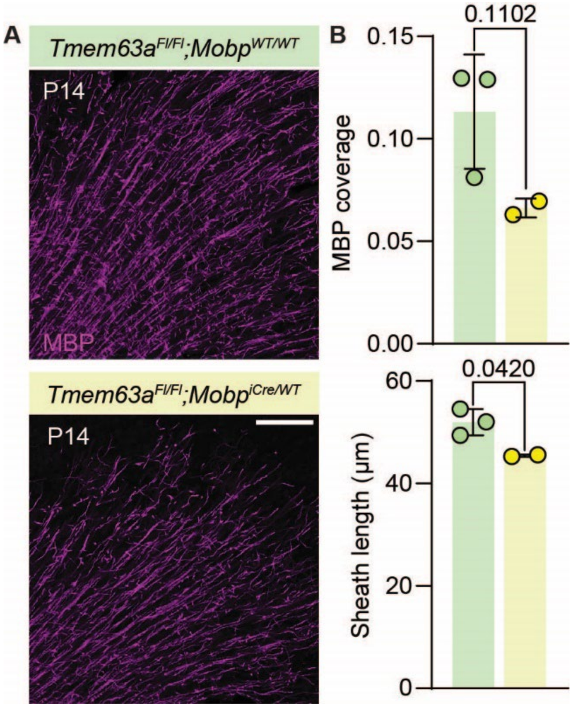


**Extended Data Fig. 3. TMEM63A functions cell autonomously in oligodendrocytes. (A)** Representative micrographs of 50 µm-thick cortical sections immunostained against MBP (magenta) from P14 *Tmem63a*$^{Fl/Fl}$*; Mobp*$^{WT/WT}$ and *Tmem63a*$^{Fl/Fl}$*; Mobp*$^{iCre/WT}$ mice. Scale bar: 100 µm. **(B)** Top, MBP coverage in cortical sections, as determined by MBP+ area divided by total imaged area, determined P14 *Tmem63a*$^{Fl/Fl}$*; Mobp*$^{WT/WT}$ and *Tmem63a*$^{Fl/Fl}$*; Mobp*$^{iCre/WT}$ mice (n = 2-3 animals, unpaired t-test). Bottom, Internode myelin sheath lengths for P14 *Tmem63a*$^{Fl/Fl}$*; Mobp*$^{WT/WT}$ and *Tmem63a*$^{Fl/Fl}$*; Mobp*$^{iCre/WT}$ mice (n = 2- 3 animals, unpaired t-test).
